# Region-specific Bmal1 deletion in the dorsal striatum alters alcohol consumption in a sex-specific manner

**DOI:** 10.64898/2026.08.02.742221

**Authors:** Mahgol Darvishmolla, Richard Courtemanche, Shimon Amir

## Abstract

**Background:** Circadian disruption is strongly associated with alcohol use disorder (AUD), but insight into the underlying brain-region and sex-specific mechanisms is limited. The function of the circadian clock gene *Bmal1* within the striatum has been linked to alcohol drinking, yet its role within functionally distinct striatal subregions has not been systematically examined.

**Methods:** We deleted *Bmal1* in medium spiny neurons of the dorsomedial striatum (DMS) or dorsolateral striatum (DLS). Male and female mice were tested for anxiety-like behavior, depressive-like behavior, and motor coordination. Voluntary alcohol intake was measured with an intermittent two-bottle choice paradigm, followed by sucrose preference and quinine-adulterated alcohol tests. To assess hormonal contributions, a subset of female mice underwent ovariectomy before behavioral testing.

**Results:** Deletion of *Bmal1* in the DLS did not alter alcohol intake, alcohol preference, or quinine-adulterated alcohol intake in either sex. In contrast, DMS *Bmal1* deletion significantly reduced alcohol consumption and alcohol preference in female mice, with no effect in males. These effects were not accompanied by changes in depressive-like behavior or motor coordination and were not explained by generalized reward changes, as sucrose preference was unaffected. Ovariectomy eliminated the effect of DMS *Bmal1* deletion on alcohol intake, indicating dependence on ovarian hormones.

**Conclusions:** The DMS is a critical site at which *Bmal1* regulates alcohol consumption in a sex-specific manner. These findings support an interaction between local circadian mechanisms and ovarian hormones in controlling alcohol drinking and highlight a potential target for sex-specific therapeutics in AUD.

## Introduction

Alcohol use disorder (AUD) is closely linked to circadian disruption, sleep disturbance, and psychiatric comorbidity (Partonen, 2015; Zhang and Volkow, 2023). At the molecular level, circadian timing is generated by transcriptional-translational feedback loops in which BMAL1 and CLOCK drive expression of Per and Cry, whose protein products then repress their own transcription (Hastings et al., 2018). Although the suprachiasmatic nucleus is the master circadian pacemaker, extra-SCN oscillators are also present in reward-related brain regions, including the striatum (Begemann et al., 2020).

The striatum is central to reward-guided action, habit formation, and addiction. Within the dorsal striatum, the dorsomedial striatum (DMS) is more strongly associated with goal-directed behavior, whereas the dorsolateral striatum (DLS) is more strongly associated with habitual responding (Corbit et al., 2012; Lipton et al., 2019; Lee et al., 2023). Because alcohol seeking is thought to shift from goal-directed to habitual control over time, these subregions are strong candidates for mediating region-specific clock-gene effects on drinking behavior.

Human and rodent studies implicate clock genes in alcohol-related phenotypes. *Bmal1* polymorphisms have been associated with alcohol use in humans (Kovanen et al., 2010; Banach et al., 2018), and manipulations of Per2, Clock, and related genes alter alcohol intake in rodents (Spanagel et al., 2005; Ozburn et al., 2013). Importantly, deletion of *Bmal1* in the whole striatum increases alcohol intake in males but suppresses it in females (de Zavalia et al., 2021), whereas deletion in the nucleus accumbens increases alcohol drinking in both sexes (Herrera et al., 2023). These findings suggest that *Bmal1* acts in a subregion-specific and sex-dependent manner within striatal circuits.

Clock genes have also been linked to affective and motivational behaviors (Dzirasa et al., 2010; Parekh et al., 2018; Schoettner et al., 2022). Because anxiety-like or depressive-like states can influence alcohol drinking, it is important to determine whether any effect of dorsal striatal *Bmal1* deletion on alcohol intake reflects a primary change in drinking behavior or a secondary change in affective state. Moreover, given established sex differences in alcohol vulnerability and striatal physiology (Flores-Bonilla and Richardson, 2020; Rangel-Barajas et al., 2021), sex must be treated as a biological variable.

Here, we tested whether selective deletion of *Bmal1* in DMS or DLS medium spiny neurons alters alcohol consumption in male and female mice. We also assessed anxiety-like behavior, depressive-like behavior, and motor coordination to evaluate behavioral specificity, and we used ovariectomy to examine the contribution of ovarian hormones to the female-specific phenotype.

## Materials and Methods

### Animals and housing

Male and female C57BL/6J mice (8–12 weeks old) carrying floxed alleles of *Bmal1* (*Bmal1*fl/fl; Jackson Laboratory, stock no. 007668) were used. Mice were housed individually in standard cages under a 12 h light/12 h dark cycle with food and water available ad libitum. Room temperature and humidity were maintained at 21 ± 1°C and 65 ± 5%, respectively. All procedures were approved by the Concordia University Animal Care Committee and complied with Canadian Council on Animal Care guidelines (certificate no. 30000256).

### Stereotaxic surgery and viral delivery

Mice underwent bilateral stereotaxic injections under isoflurane anesthesia using a SomnoSuite low-flow vaporizer system. Anesthesia was induced at 3–4% isoflurane in oxygen and maintained at 1–2%. Carprofen (5 mg/kg) was administered as perioperative analgesia.

To delete *Bmal1* selectively in neurons, *Bmal1*-floxed mice received bilateral injections of AAV2/5-hSyn-CRE-EGFP (1.0 × 10^12 vg/ml), whereas controls received AAV2/5-hSyn-EGFP (1.0 × 10^12 vg/ml). Target coordinates were as follows: DMS, AP +0.98 mm, ML ±1.2 mm, DV −2.85 mm; DLS, AP +0.98 mm, ML ±2.2 mm, DV −2.8 mm. Virus was delivered bilaterally at 400 nl/min. The hSyn promoter restricted transgene expression to neurons. At the end of the study, brains were examined by fluorescence microscopy to verify infection. Mice were excluded if expression was unilateral or insufficient in localization or spread.

### Ovariectomy

A subset of female mice underwent bilateral ovariectomy under isoflurane anesthesia. After a dorsal midline incision, the ovaries were exteriorized through small muscle-wall openings, the oviducts were ligated, and the ovaries were removed, as described previously (Souza et al., 2019).

### Behavioral procedures

Behavioral testing was performed after a 3-week postsurgical recovery period and proceeded in order of increasing stress, with at least 1 week between tests (Fig. 1). Animals were habituated to the testing room for 1 h before testing. Most procedures were conducted between ZT2 and ZT6; Rotarod testing was conducted between ZT6 and ZT8, and the tail suspension test at ZT8.

**Fig. 1.**
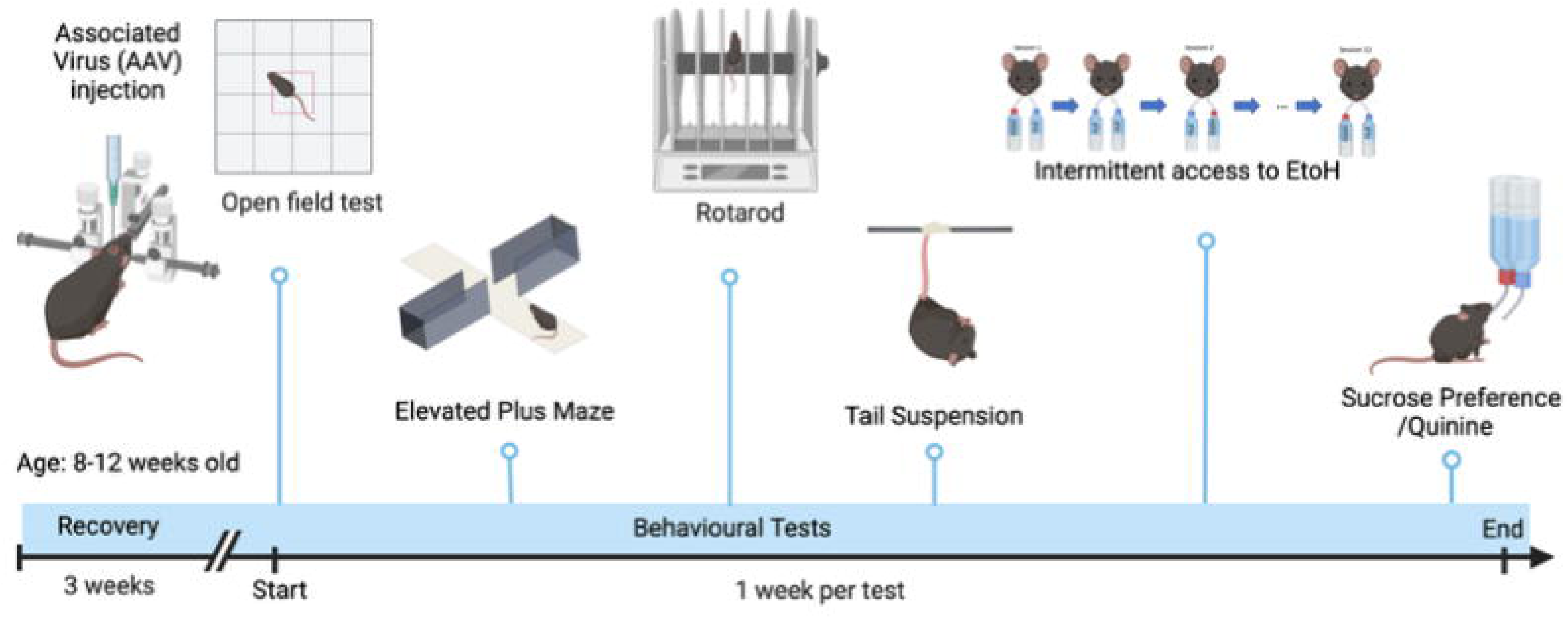
Experimental timeline. Following stereotaxic surgery, mice recovered for 3 weeks before behavioral testing, which was conducted in order of increasing stress, with approximately 1 week between assays.

### Elevated plus maze

The elevated plus maze (EPM) was used to assess anxiety-like behavior (Walf and Frye, 2007). The apparatus consisted of two open arms and two closed arms connected by a center platform. Mice were placed in the center facing a closed arm, and behavior was video recorded. Time spent in the open arms and number of open-arm entries were quantified.

### Open field test

The open field test (OFT) was used to assess anxiety-like behavior based on avoidance of the exposed center (Seibenhener and Wooten, 2015). Mice were placed in an open arena equipped with infrared beams and tracked for 30 min. Total distance traveled, time spent in the center, and latency to enter the center were measured.

### Rotarod

Motor coordination was assessed with a fixed-speed rotarod, given the DLS contribution to sensorimotor processing (Lipton et al., 2019). Mice first underwent a habituation session at 2 rpm. During testing, animals completed three 1-min trials each at 4, 8, and 12 rpm, with 30 s rest intervals. Mice unable to complete 8-rpm trials were not tested at 12 rpm. Performance was expressed as cumulative time on the rod.

### Tail suspension test

The tail suspension test (TST) was used to assess depressive-like behavior (Can et al., 2012). Mice were suspended by the tail for 6 min, and immobility time was calculated from video recordings as total suspension time minus mobility time.

### Intermittent access to ethanol

Voluntary alcohol intake was measured using an intermittent two-bottle choice procedure. After 3 habituation days with water in both bottles, mice underwent twelve 24-h sessions with access to water and 15% (v/v) ethanol. Bottle positions were alternated to control for side preference. Ethanol intake was expressed as g/kg/session, and alcohol preference was calculated as ethanol intake divided by total fluid intake.

### Sucrose preference and quinine-adulterated alcohol

To assess generalized reward processing, mice underwent a 3-day sucrose preference test with access to water and 1% sucrose. To assess aversion-resistant alcohol drinking, mice were then tested for 3 sessions with access to water and ethanol containing 0.25% quinine. Sucrose preference and quinine-adulterated alcohol intake were quantified from bottle weights.

### Western Blot

Striatal punches were lysed in NP40 lysis buffer containing 1% Nonidet P40, 0.1% SDS, 50 mM Tris base, 0.1 mM EDTA, 0.1 mM EGTA, and 0.1% deoxycholic acid, pH 7.4. The buffer containing inhibitor cocktails composed of 1 mM sodium pyrophosphate, 1 mM sodium orthovanadate, 20 mM NaF, a protease inhibitor, and a phosphatase inhibitor (Sigma-Aldrich, P5726-1ML) at 1 mM was added to the NP40 buffer. The lysate was incubated under rotation for 30 minutes at 4°C. Proteins containing supernatant were collected after centrifugation at 14,000 rpm for 10 minutes at 4°C. Protein quantification was performed using the Pierce BCA Protein Assay Kit (ThermoScientific, #23227) with a BSA standard curve from 0 to 1 μg/μL. An equal amount of protein 30 μg per sample, was mixed with 4X Laemmli loading buffer (Bio-Rad,#1610747) in a total volume of 60 μL and heated to 95°C for 5 minutes to denature the proteins. Samples were separated by SDS-PAGE on a 10% acrylamide gel using the Mini-PROTEAN Tetra System (Bio-Rad). Electrophoresis was performed in 1X Tris-Glycine electrophoresis buffer for 30 minutes at 100 V to gather the proteins, then one hour at 120 V for migration in the separation gel. The transfer was done onto a 0.2 μm nitrocellulose membranev(Bio-Rad, #1620112) for one hour at 100V in cold 1X Tris-Glycine-methanol transfer buffer. Membranes were blocked for 1 hour at room temperature in 5% TBST-BSA according to the manufacturer’s recommendations. Blocked membranes were then incubated at 4°C with the primary antibody (2.5% TBST-BSA) until the next day before being incubated with the secondary antibody (1/3000) of rabbit or mouse (Goat Anti-Rabbit IgG (H+L) HRP Conjugate, Millipore, LV1646281, Goat Anti-Mouse IgG (H+L) HRP Conjugate, Millipore, AP124P) for one hour at room temperature. Proteins were detected using the Clarity Western ECL Substrate 36 detection system (Bio-Rad, #170-5061). Primary antibodies used are: Bmal1 (Novus Biologicals, NB100-2288, Dilution 1:1000, Rabbit) and β-actin (Sigma, A5316, Dilution 1: 2000, Mouse). Densitometric analyses of immunoblots were performed using ImageJ software (Fiji, Version 2.1.0/1.53c).

### Statistics

Data were analyzed in GraphPad Prism 9 and are presented as mean ± SEM. Behavioral and alcohol-drinking data were analyzed with two-way ANOVA using sex and viral vector as factors, followed when appropriate by Šídák-corrected post hoc comparisons. Sucrose preference and quinine-adulterated alcohol measures were analyzed with unpaired two-tailed t tests when normality assumptions were met; otherwise, Mann–Whitney tests were used. Statistical significance was set at p ≤ 0.05.

## Results

### Validation of Bmal1 deletion

At the conclusion of the experiments, viral transduction was confirmed histologically. GFP and Cre expression were observed in medium spiny neurons within the targeted DMS or DLS, and representative immunofluorescence images confirmed regional *Bmal1* deletion in animals (Fig. 2A & B).

**Fig. 2.**
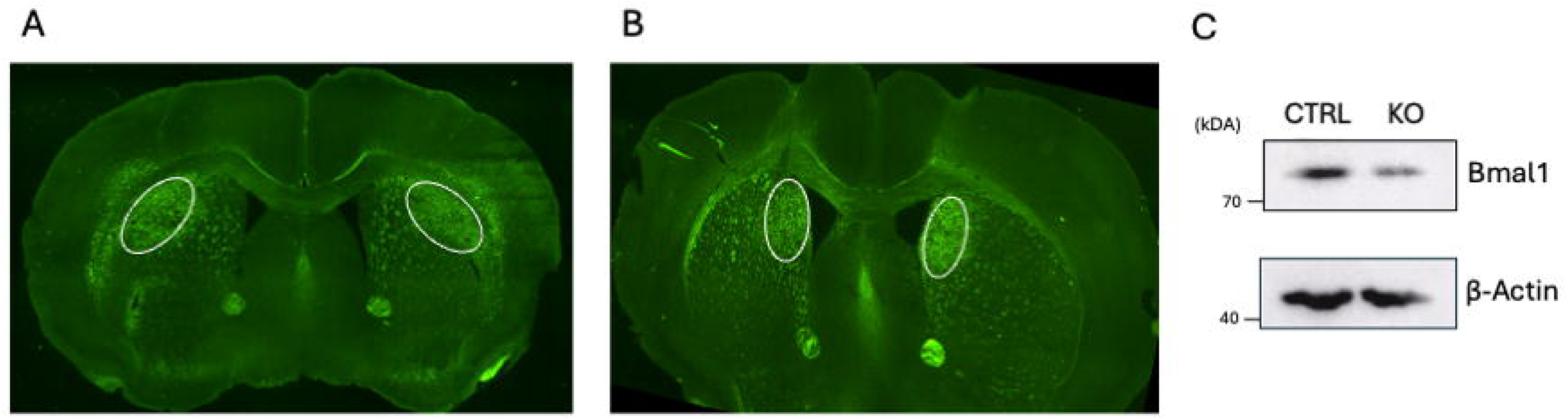
Representative verification of viral targeting in DLS and DMS after injection of AAV2/5-hSyn-CRE-EGFP. A & B represent the strong EGFP expression in the DLS and DMS region, respectively, indicating successful viral transduction and Cre-recombinase expression in the targeted regions, mediating the site-specific deletion of the *Bmal1.* Western blotting reports the efficacy of the AAV2/5-hSyn-CRE-EGFP to delete BMAL1 in the striatum tissue of KO and Ctrl animals.

Western blot analysis reported the deletion of *Bmal1* from the DMS and DLS collected from animals infused with a virus expressing Cre protein, and these were compared to controls (Fig. 2C).

### DLS Bmal1 deletion and behavior

Deletion of *Bmal1* in DLS medium spiny neurons did not alter most measures of anxiety-like behavior, motor coordination, or depressive-like behavior (Fig. 3). In the EPM, total distance traveled [F(1,17)=0.9005, p=0.3559] and open-arm entries [F(1,17)=0.03817, p=0.8474] were unchanged. However, time spent in the open arms was increased in knockout mice of both sexes [F(1,17)=13.40, p=0.0019], suggesting an anxiolytic-like effect in this assay. In the OFT, total distance traveled [F(1,17)=0.6022, p=0.4484], time spent in the center [F(1,17)=1.058, p=0.3180], and latency to enter the center [F(1,17)=1.067, p=0.3161] were unaffected. Rotarod performance [F(1,17)=0.07693, p=0.7849] and TST immobility [F(1,17)=0.2642, p=0.6139] were also unchanged.

**Fig. 3.**
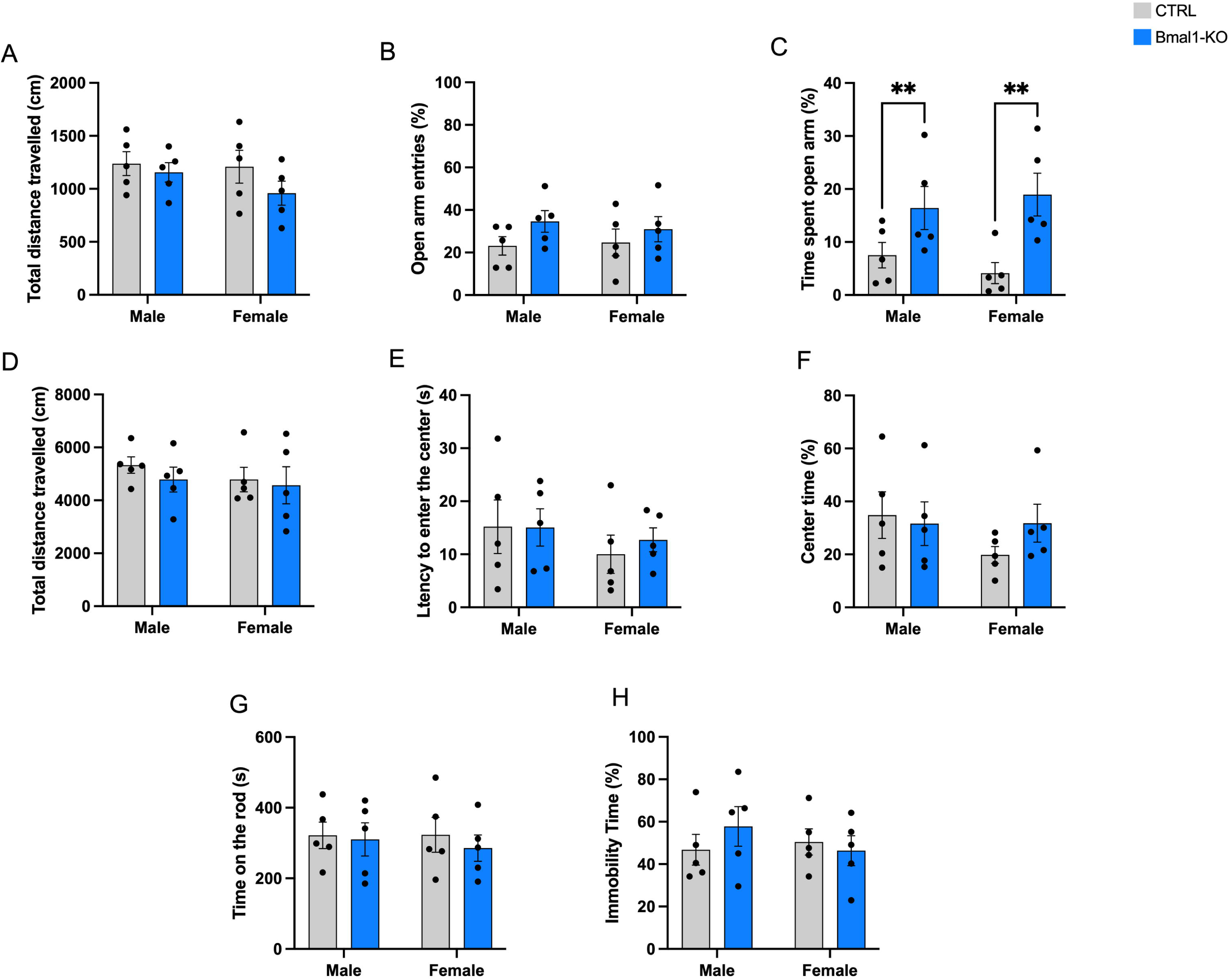
Effects of DLS *Bmal1* deletion on anxiety-like behavior, motor coordination, and depressive-like behavior. EPM (A–C), OFT (D–F), Rotarod (G), and TST (H). DLS knockout increased time in the open arms of the EPM but did not alter other behavioral measures. (n=5 among groups and sexes).

### DLS Bmal1 deletion and alcohol-related measures

DLS *Bmal1* deletion did not affect alcohol drinking in either sex (Fig. 4). Across 12 intermittent-access sessions, alcohol intake was unchanged in females [F(1,84)=0.3146, p=0.5764] and males [F(1,7)=3.551, p=0.1015]. Alcohol preference was similarly unaffected in females [F(1,8)=4.508, p=0.0665] and males [F(1,8)=0.4679, p=0.5133]. Sucrose preference did not differ between knockout and control mice in males (p=0.2207) or females (p=0.6643). Quinine-adulterated alcohol intake was also unchanged in males (p=0.5750) and females (p=0.0528). Together, these results indicate that *Bmal1* in the DLS does not measurably regulate voluntary alcohol intake under these conditions.

**Fig. 4.**
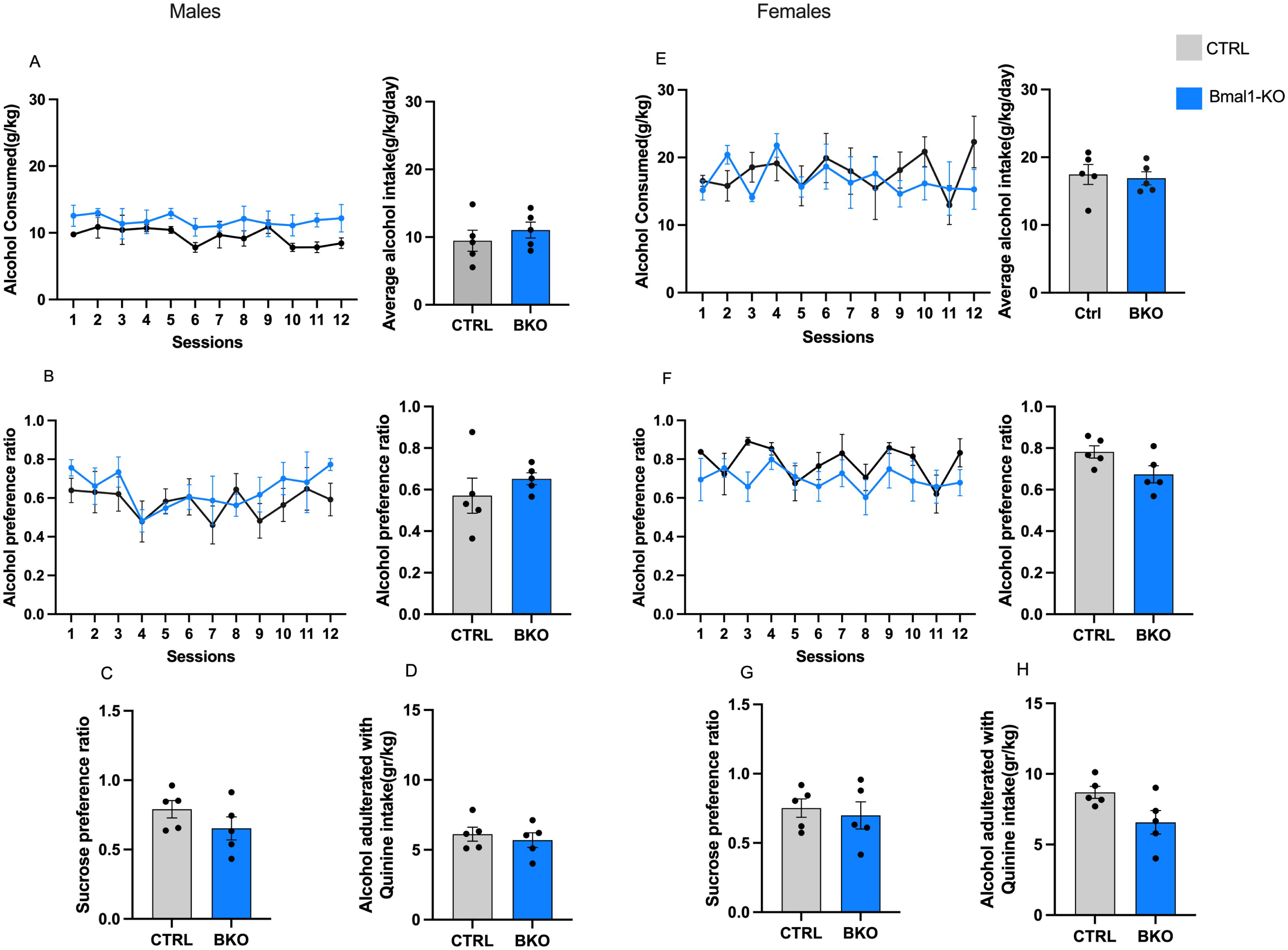
Alcohol-related measures after DLS *Bmal1* deletion. Daily and average alcohol intake and preference, sucrose preference, and quinine-adulterated alcohol intake were not altered in either sex. (n=5 among groups and sexes).

### DMS Bmal1 deletion and behavior

In the DMS, *Bmal1* deletion did not broadly alter anxiety-like, depressive-like, or motor behaviors (Fig. 5). In the EPM, total distance traveled [F(1,28)=2.223, p=0.1472], time spent in the open arms [F(1,28)=0.05292, p=0.8197], and open-arm entries [F(1,28)=5.705, p=0.0239] were unaffected. In the OFT, total distance traveled [F(1,28)=0.03445, p=0.8541] and latency to enter the center [F(1,28)=3.7, p=0.0620] were unchanged. However, female knockout mice spent more time in the center than male knockout mice [F(1,28)=4.680, p=0.0392], consistent with a modest sex-dependent anxiolytic-like effect. Rotarod performance [F(1,28)=1.862, p=0.1834] and TST immobility were not affected [F(1,28)=0.3358, p=0.5669].

**Fig. 5.**
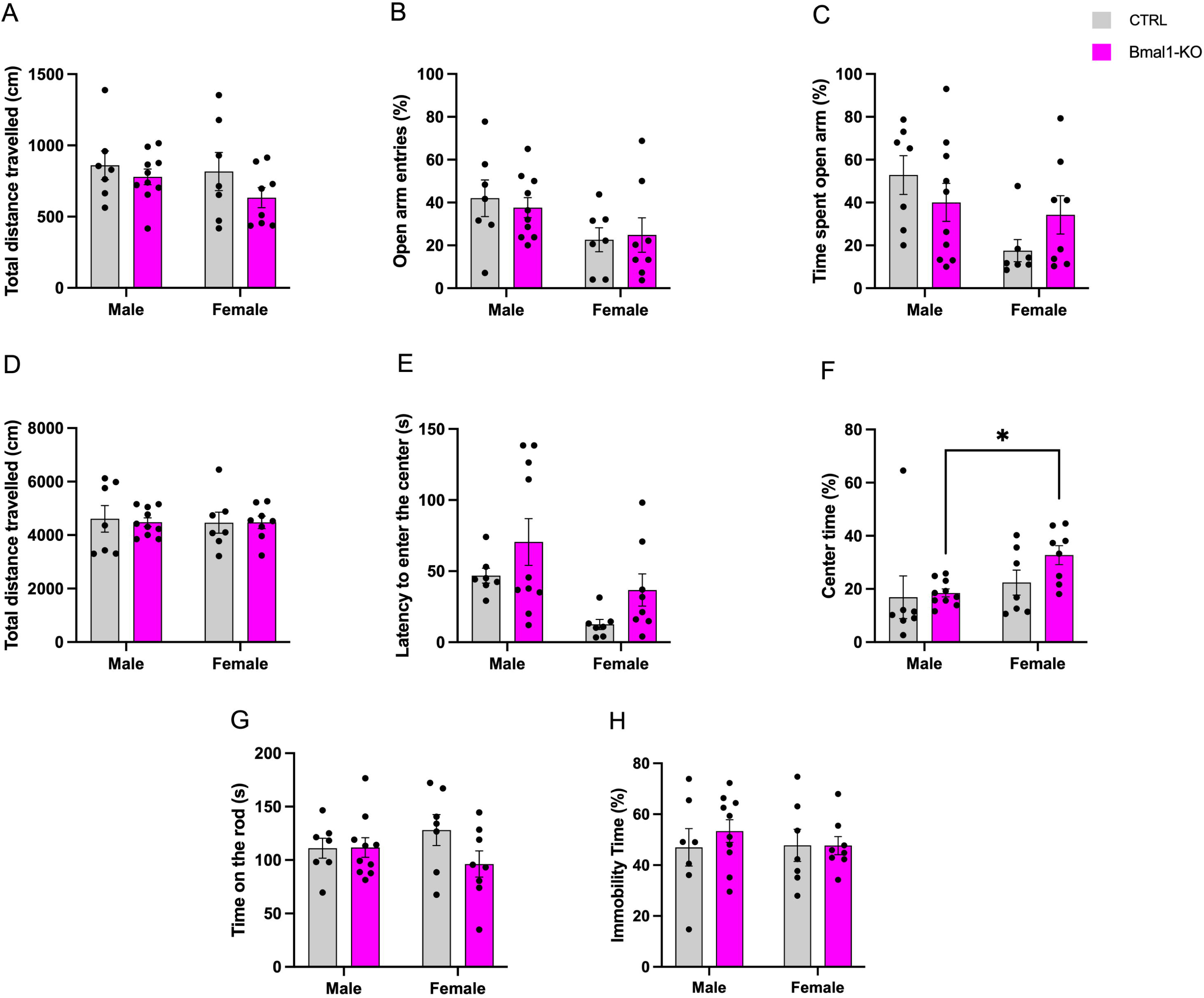
Effects of DMS *Bmal1* deletion on anxiety-like behavior, motor coordination, and depressive-like behavior. EPM (A–C), OFT (D–F), Rotarod (G), and TST (H). Most measures were unchanged, although female knockout mice spent more time in the OFT center than male knockout mice. Female CTRL group (n=7), male CTRL (n=7), male *Bmal1-*KO group (n=10), and the female *Bmal1-*KO group (n=8).

### DMS Bmal1 deletion and alcohol-related measures

By contrast, DMS *Bmal1* deletion altered alcohol drinking in a sex-specific manner (Fig. 6). In males, knockout mice did not differ from controls in alcohol intake [F(1,15)=2.587, p=0.1286] or alcohol preference [F(1,15)=1.505, p=0.2389]. In females, however, knockout mice showed reduced alcohol intake [F(1,13)=15.04, p=0.0019] and reduced alcohol preference [F(11,143)=2.404, p=0.0091] relative to controls. Average alcohol intake was reduced by 20.6%, and female knockout mice consumed an average of 8.24 g/kg/session less alcohol than controls. Alcohol preference was reduced by approximately 25% across the 12 drinking sessions.

**Fig. 6.**
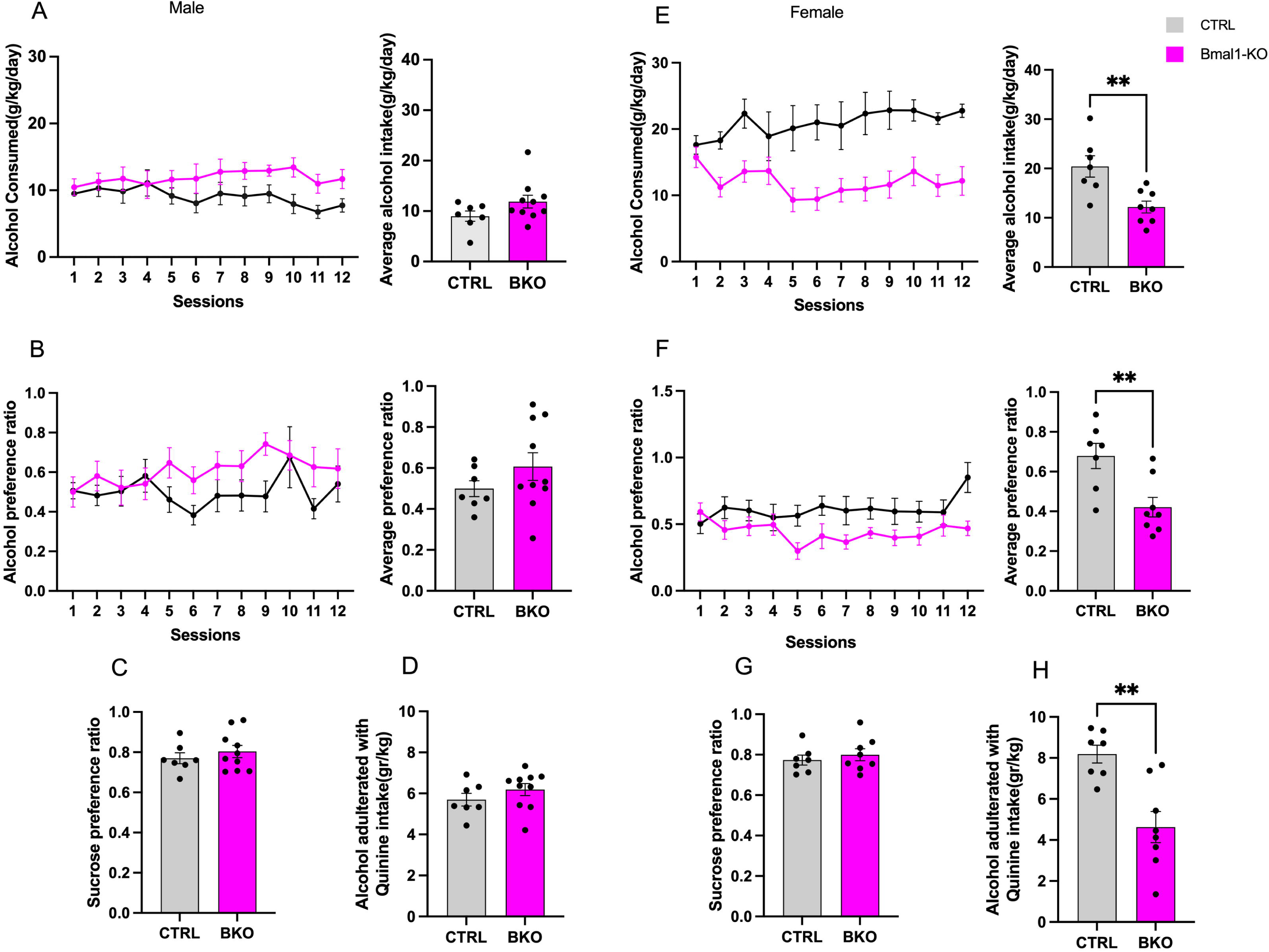
Alcohol-related measures after DMS *Bmal1* deletion. In males, alcohol intake and preference were unchanged. In females, DMS knockout reduced alcohol intake, alcohol preference, and quinine-adulterated alcohol intake without altering sucrose preference. Female CTRL group (n=7), male CTRL group (n=7), *male Bmal1-*KO (n=10), and female *Bmal1*-KO (n=8).

These effects were selective for alcohol drinking: sucrose preference was unchanged in females (p=0.5501) and males (p=0.4946), arguing against a generalized reward deficit. In the quinine test, female knockout mice consumed less quinine-adulterated alcohol than female controls (p=0.0018), whereas males showed no difference (p=0.2784), indicating that DMS *Bmal1* downregulation in females also reduces intake of aversive alcohol solutions.

### Ovariectomy

To assess the contribution of ovarian hormones, we compared ovariectomized female DMS knockout mice (n=6) with ovariectomized controls (n=5) (Fig. 7). No significant group differences were observed in alcohol intake [F(1,9)=3.615, p=0.0897], alcohol preference [F(1,9)=1.194, p=0.3029], sucrose preference (p=0.8575), or quinine-adulterated alcohol intake (p=0.3579). Notably, ovariectomized knockout mice showed 20.96% greater alcohol intake than intact knockout females. These findings indicate that the suppressive effect of DMS *Bmal1* deletion on alcohol drinking depends on ovarian hormones.

**Fig. 7.**
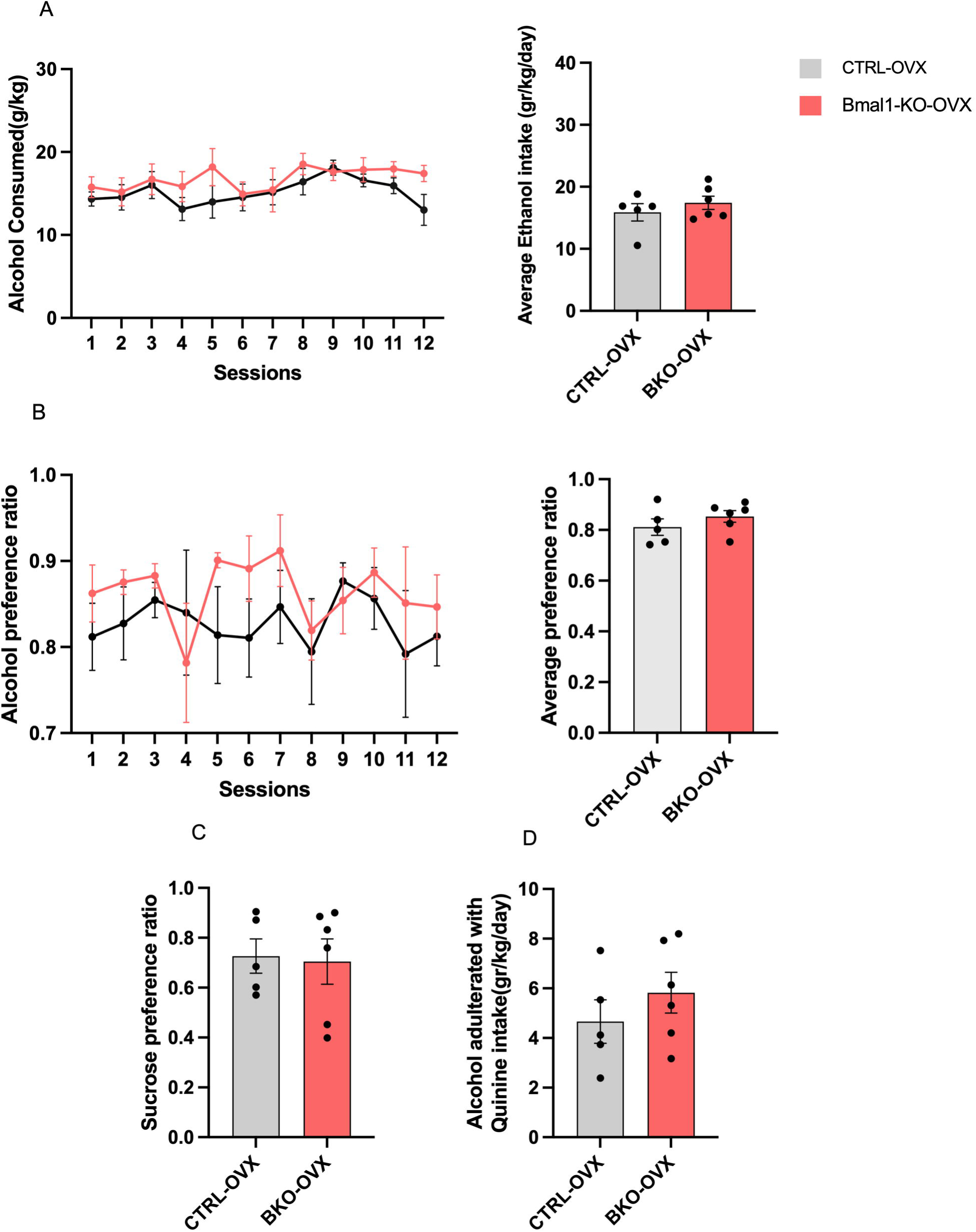
Alcohol-related measures in ovariectomized female mice with DMS *Bmal1* deletion. Alcohol intake, alcohol preference, sucrose preference, and quinine-adulterated alcohol intake did not differ between ovariectomized knockout and ovariectomized control groups. BKO-OVX mice (n=6) and CTRL-OVX group (n=5)

## Discussion

This study demonstrates that *Bmal1* regulates alcohol consumption in a region- and sex-specific manner within the dorsal striatum. Deletion of *Bmal1* in DMS medium spiny neurons reduced voluntary alcohol intake and preference in females but had no effect in males, whereas deletion in the DLS had no effect on alcohol drinking in either sex. The female-specific DMS phenotype was abolished by ovariectomy, indicating dependence on ovarian hormones. Behavioral testing further showed that these drinking effects were not accompanied by changes in depressive-like behavior or motor coordination and were not explained by altered sucrose preference.

The regional dissociation between DMS and DLS is consistent with known functional specialization of dorsal striatal circuits. The DMS is more strongly involved in goal-directed behavior, whereas the DLS is more strongly linked to habit formation (Corbit et al., 2012; Lipton et al., 2019). Because the intermittent-access procedure used here is not designed to induce long-term habit-like alcohol seeking, it is plausible that DMS circuits would have a greater influence than DLS circuits on the measured phenotype. In this context, *Bmal1* may modulate alcohol drinking through neural processes more closely tied to action valuation and decision-making than to established habitual responding.

The present findings also refine prior work showing that whole-striatum *Bmal1* deletion decreases alcohol consumption in females but increases it in males (de Zavalia et al., 2021), and that nucleus accumbens *Bmal1* deletion increases alcohol drinking in both sexes (Herrera et al., 2023). Together, these studies support the view that striatal *Bmal1* does not exert a uniform effect. Instead, the behavioral outcome depends on the subregion targeted and on sex, with DMS deletion producing an intake-suppressing effect in females that appears to dominate the whole-striatum phenotype.

Several mechanisms may underlie this effect. Clock genes influence glutamatergic, GABAergic, and dopaminergic signaling, all of which are central to dorsal striatal function. Chronic alcohol exposure alters prelimbic cortical output to the DMS and disrupts genes involved in synaptic transmission, including Syt1 (Tapocik et al., 2013; Barbier et al., 2015, 2021). Other clock-gene models have shown altered AMPA receptor regulation and GABA_a_ receptor subunit expression (Dzirasa et al., 2010; Parekh et al., 2018; Ozburn et al., 2017). Dopamine signaling is also tightly coupled to circadian mechanisms (Hood et al., 2010; Kiehn et al., 2023), and alcohol exposure alters dorsal striatal dopamine release in a sex-dependent manner (Salinas et al., 2021; Kania et al., 2025). Although the present study was not designed to identify a cellular mechanism, the data are consistent with the possibility that loss of *Bmal1* changes DMS circuit function by altering excitatory, inhibitory, and/or dopaminergic transmission.

The ovarian hormone dependence of the female phenotype is especially notable. Estradiol receptors are highly expressed in the dorsal striatum, estradiol modulates dorsal striatal plasticity and dopamine signaling, and ovarian hormones promote alcohol intake in female mice (Ramôa et al., 2013; Satta et al., 2018; Lewitus and Blackwell, 2023; Lewitus et al., 2024). The absence of a knockout effect after ovariectomy therefore suggests that *Bmal1* acts within a hormone-sensitive DMS network. One plausible interpretation is that ovarian hormones enable or amplify the circuit consequences of *Bmal1* loss, thereby revealing a female-specific suppression of alcohol drinking.

Some behavioral effects outside alcohol intake were observed, but they were limited. DLS knockout mice spent more time in the open arms of the EPM, and female DMS knockout mice spent more time in the center of the OFT than male knockouts, suggesting modest anxiolytic-like effects in selected assays. However, the absence of broad effects across anxiety, depressive-like behavior, sucrose preference, and motor coordination indicates that the principal phenotype is selective to alcohol-related behavior rather than a general change in locomotion, motivation, or reward sensitivity.

This study has several limitations. First, the viral strategy targeted medium spiny neurons broadly and did not distinguish between D1- and D2-expressing populations. Second, the mechanistic conclusions are necessarily indirect because receptor expression, synaptic physiology, and local circuit dynamics were not measured.

Third, the DLS-negative result may depend in part on the behavioral model used, which emphasizes voluntary drinking rather than established habit. Future studies incorporating cell-type-specific targeting, electrophysiology, and extended drinking paradigms will be important for defining the underlying circuit mechanisms more precisely.

In summary, these findings identify the DMS as a key locus at which *Bmal1* regulates alcohol consumption in a sex-specific manner. The results further indicate that ovarian hormones are required for the suppressive effect of DMS *Bmal1* deletion on alcohol intake in females. This work advances understanding of how local circadian mechanisms interact with sex-dependent striatal circuitry to shape alcohol-related behavior and may inform the development of sex-specific approaches to AUD treatment.

## Acknowledgements

We wish to thank Konrad Schöttner for his invaluable support and coordination of multiple laboratory operations. We also thank Amanda Szubinski for her technical help.

## Author contributions

Conceptualization, data curation: MD & SA; supervision, validation, funding acquisition & project administration: RC & SA; writing, original draft: MD; formal methodology and analysis, writing – review and editing: MD, SA & RC.

## Ethical approvals

This project was approved by the Concordia University Animal Research Ethics Committee, following the policies from the Canadian Council on Animal Care.

